# A mitochondrial sirtuin shapes the intestinal microbiota by controlling lysozyme expression

**DOI:** 10.1101/2023.06.02.543385

**Authors:** Mirjam Knop, Christian Treitz, Stina Bettendorf, Judith Bossen, Jakob von Frieling, Shauni Doms, Iris Bruchhaus, Ronald P. Kühnlein, John F. Baines, Andreas Tholey, Thomas Roeder

**Author notes:** correspondence: Thomas Roeder, CAU Kiel, Zoology, Olshausenstrasse 40, 24098 Kiel.

## Abstract

Sirtuins act as cellular sensors in the gut that control a substantial change in gut properties in response to environmental changes. Here we show that the only mitochondrial sirtuin of *Drosophila, dSirt4*, is strongly up-regulated by a protein-reduced diet. Flies with a dSirt4 defect show strong changes in the protein pattern and physiological properties of their intestine. One of the most notable effects was the strong induction of lysozyme gene expression in the intestine, which also translates into enhanced lysozyme activity. This effect was cell autonomous, as it was also observed in flies with *dsirt4* was exclusively silenced in enterocytes of the intestine. Although this strongly increased lysozyme expression, it did not reduce total bacterial load in the intestine, but rather changed the composition of the microbiota by reducing the number of gram-positive bacteria. This effect on microbiota composition can be attributed to the dSirt4-dependent lysozyme expression, as it was absent in a lysozyme-deficient background. *dSirt4* deficiency in enterocytes reduced lifespan of flies, which was also observed in those flies experiencing ectopic lysozyme overexpression in enterocytes. This implies that strong lysozyme expression leads to a dysbiotic state associated with reduced lifespan.

## Introduction

The intestinal epithelium is a central metabolic organ responsible for orchestrating a wide range of metabolic aspects of the organism. Despite its seemingly simple architecture, intestinal epithelia are characterized by a marked cellular complexity. This complexity appears to be phylogenetically conserved as it is observed throughout the animal kingdom (1, 2). Together with the endogenous microbiota, the intestinal epithelium is constantly exposed to a wide range of environmental factors (3, 4). The most important of these environmental factors is the diet, which is highly variable in terms of energy content and quality (5). As a result, one of the main functions of the intestinal epithelium is the maintenance of homeostasis under these changing environmental conditions. This maintenance of homeostasis involves not only the intestinal epithelium itself but also the closely associated microbiota (6). Disruption of this homeostatic situation is causally linked with microbiota dysbiosis and intestinal diseases, including inflammatory bowel diseases (IBDs) (7, 8).

Proteins that combine an energy-sensing function with the potential for long-term and widespread adaptation of cellular metabolic properties are particularly suited for this adaptation. Sirtuins, nicotinamide adenine dinucleotide (NAD^+^)-dependent protein deacetylases, can fulfill this role (9). Sirtuins are a group of highly conserved proteins sharing homology with the yeast silent information regulator 2 (Sir2) (10) that are involved in a wide range of biological processes, including metabolism, aging, DNA repair, and regulation of the microbiota (11, 12). They are operative in all organs including the intestine where they help to maintain the functional and structural integrity of the intestine (13). The observation that even in planarians sirtuin activity is required to maintain intestinal integrity implies a phylogenetically ancient role of this protein family in keeping organ homeostasis (14). Studies using mice deficient for Sirt1 only in enterocytes of the gut showed that this sirtuin is required to prevent the development of IBD. This effect seems to be mediated via shaping the gut microbiota (15). Therefore, Sirt1 activity is needed to prevent the translocation of harmful bacteria from the intestinal lumen into the bloodstream (15). Sirt1-deficient intestines have more goblet and Paneth cells, thus changing the antimicrobial tone in the intestinal lumen, which in turn can affect the composition of the gut microbiota (16). Moreover, reduced Sirt1 levels are found in biopsies of IBD patients (17) and Sirt1 is required for bile acid absorption through directly targeting the HNF1α/FXR signaling (18). Besides the effects of Sirt1 on the composition of the microbiota, especially beneficial members of the microbiota can also upregulate the intestinal expression of Sirt1 (19), therewith showing that the interaction between sirtuin signaling and the microbiota is highly interconnected.

A subgroup within the sirtuin family, the mitochondrial sirtuins, which include sirtuins 3, 4 and 5, have attracted particular interest because they act directly in the energy centers of the cell (20, 21). This important role in metabolic control resulting from this activity in mitochondria is also particularly important because IBDs are associated with mitochondrial dysfunction (22, 23). As an example, highlighting the important role of mitochondrial sirtuins in maintaining intestinal homeostasis, it was shown that dysfunctional SIRT3 expression comes along with impaired gut barrier function, which in turn results from a dysbiotic microbiota (24). Despite these studies showing a key role for sirtuins, especially mitochondrial sirtuins, in maintaining intestinal homeostasis, our mechanistic understanding of these processes is still rudimentary.

Simpler models are helpful to better understand the general importance of mitochondrial sirtuins and their role in controlling the composition of the microbiota. *Drosophila melanogaster* contains only one mitochondrial sirtuin, dSirt4. This sirtuin has a significant influence on lifespan, and a modest fat body-targeted overexpression of this gene leads to a prolongation of life. In contrast, *dSirt4*-deficient animals have a shorter lifespan. This could be linked to altered metabolic properties. dSirt4 plays an important role in the communication between cells and their mitochondria in a genotype-specific manner (25). In addition, a first connection between dSirt4 and the microbiota could be demonstrated via the interaction between the endosymbiont *Wolbachia* and the expression level of *dSirt4* (26). To gain a deeper and more mechanistic understanding of the processes controlled by dSirt4 in the intestine, we turned to the fruit fly *Drosophila*. We found that some of the phenotypes observed in *dSirt4*-deficient animals also occur in animals in which *dSirt4* is silenced only in the enterocytes of the intestine. The most exciting finding was that *dSirt4* deficiency in enterocytes leads to a dramatic increase in lysozyme activity, which directly affects the composition of the microbiota.

## Results

To identify those sirtuins that are of particular interest for adaptation to changing environmental conditions in the gut, we performed a transcriptome analysis. To do this, we exposed adult females (*w*^1118^) to different nutritional conditions, i.e. massive food stress. We used a holidic diet as described recently (27, 28), which allowed us to keep all dietary components constant, except for the protein content of the diet, which was reduced to zero in this experiment. This feeding intervention lasted for 7 days, after which the guts of the animals were isolated and compared with age-matched controls. *dSirt4*, which encodes the single mitochondrial sirtuin of *Drosophila* was the only of the five *fly* sirtuins, significantly up-regulated by the feeding stress. In contrast, other sirtuins were down-regulated (Fig. 1A). Based on these results, all further investigations were focused on *dSirt4*.

**Fig. 1:**
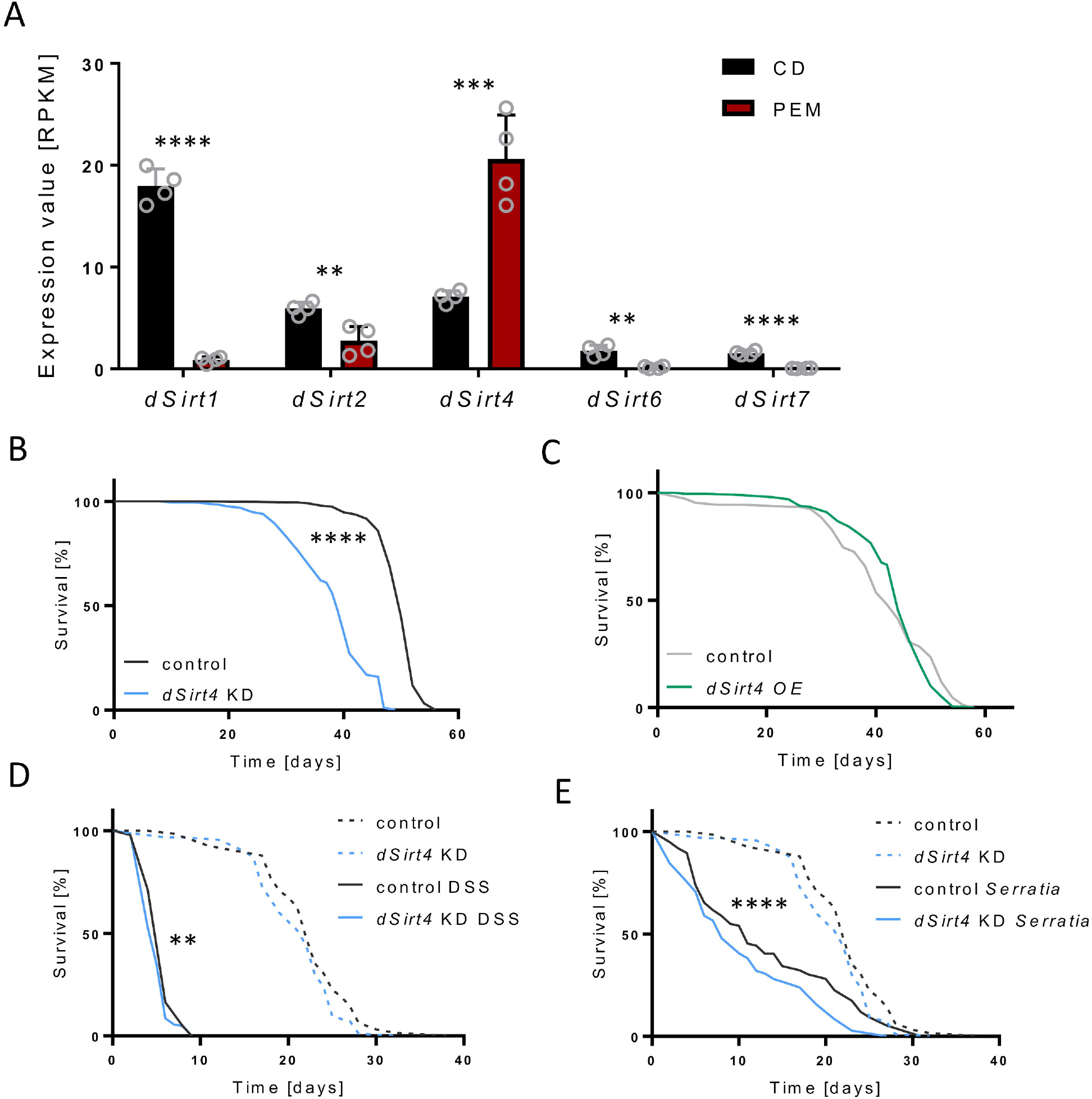
dSirt4 has an important role in the stress response of the *Drosophila* intestine. (A) A transcriptome analysis of isolated intestines of flies subjected to 7 days protein depleted diet (PEM) revealed that the only sirtuin gene upregulated under these conditions is the only mitochondrial sirtuin of *Drosophila, dSirt4*, while all others were downregulated (CD = control diet, n = 4). (B) A life span analysis showed a significantly reduced survival of flies with a knockdown of *dSirt4* specifically in enterocytes (n = 187-195). (C) The overexpression of *dSirt4* in enterocytes did not affect the life span of flies (n = 91-185). (D) *dSirt4* deficient flies showed reduced survival in response to infection with Serratia marcescens, while the control treatment with sucrose did not lead to changes in life span (n = 63-104). (E) In response to the treatment with DSS, *dSirt4* deficient flies lived shorter than their genetic control (n = 63-186). (B, D, E) control = *w*^1118^>UAS-sirt4 crispr/Cas9, *dSirt4* KD = NP1-Gal4;tubPGal80ts>UAS-sirt4 crispr/Cas9, (C) control = *w*^1118^ >UAS-sirt4, dSirt4 = NP1-Gal4;tubPGal80ts > UAS-sirt4. * = p < 0.05, ** = p < 0.01, **** = p < 0.0001.

To characterize the role of dSirt4 in the intestine, we first measured the life span of flies with a Crispr/Cas9 mediated knockdown of *dSirt4* specifically in enterocytes (*NP1-Gal4; tubPGal80ts>dsirt4-sgRNA; UAS-Cas9*). We found that these flies showed a significantly reduced survival with a median life span of 39 days compared to the effector control (*w*^1118^*>dsirt4-sgRNA; UAS-Cas9)*, which had a median life span of 50 days (Fig. 1B). However, the over-expression of *dSirt4* specifically in enterocytes (*NP1-Gal4; tubPGal80ts>UAS-dsirt4)* did not affect the survival of flies (median survival: 42-44 days, Fig. 1C). Next, we tested whether a deficiency in *dSirt4* expression in enterocytes influences the susceptibility to orally applied stressors. Here, infection with *Serratia marcescens*, an entomopathogenic gram-negative bacterium (29) was used to quantify the effects of *dSirt4*-deficiency. Flies with a knockdown of *dSirt4* in enterocytes lived significantly shorter compared to *w*^1118^ flies upon constant bacterial infection (Fig. 1D). The median life span was reduced to 8 days compared to the infected control (*w*^1118^, 11 days). Surprisingly, *dSirt4* knockdown flies in the infection-free control, where flies were fed on 5% sucrose only, showed no significant difference in survival compared to their genetic control (median life span = 22 days; Fig. 1D). Dextran Sodium Salt (DSS) is a substance used to induce colitis in mice (30), in *Drosophila*, DSS also induces tissue damage and proliferation as well as a reduction of lifespan (31). As expected, the application of 5 % DSS in 5 % sucrose solution strongly reduced the survival of flies compared to the DSS-free controls. This reduction was slightly, but significantly more pronounced in flies deficient in *dSirt4* (Fig. 1E). Compared to the genetic control, the median survival of *dSirt4* deficient flies was reduced from 6 to 5 days.

To further characterize the role of dSirt4, we analyzed the gut functionality. Therefore, we quantified the daily consumption of food (32). *dSirt4*-deficient flies consumed significantly more food compared to the control (Fig. 2A), but they surprisingly excreted fewer fecal spots (Fig. 2B). Flies with *dSirt4*-deficiency only in enterocytes showed no significant difference in the amount of ingested food compared to the control (Fig. 2C). However, the number of fecal spots was reduced (Fig. 2D). Furthermore, the metabolic rate of *dsirt4*-deficient flies was evaluated by measuring heat dissipation. Compared to the control, no significant changes were detectable (Fig. 2E). Finally, we measured the spontaneous locomotor activity of flies experiencing enterocyte-specific knockdown of *dSirt4*. Interestingly, these flies had a significantly reduced activity (Fig. 2F).

**Fig. 2:**
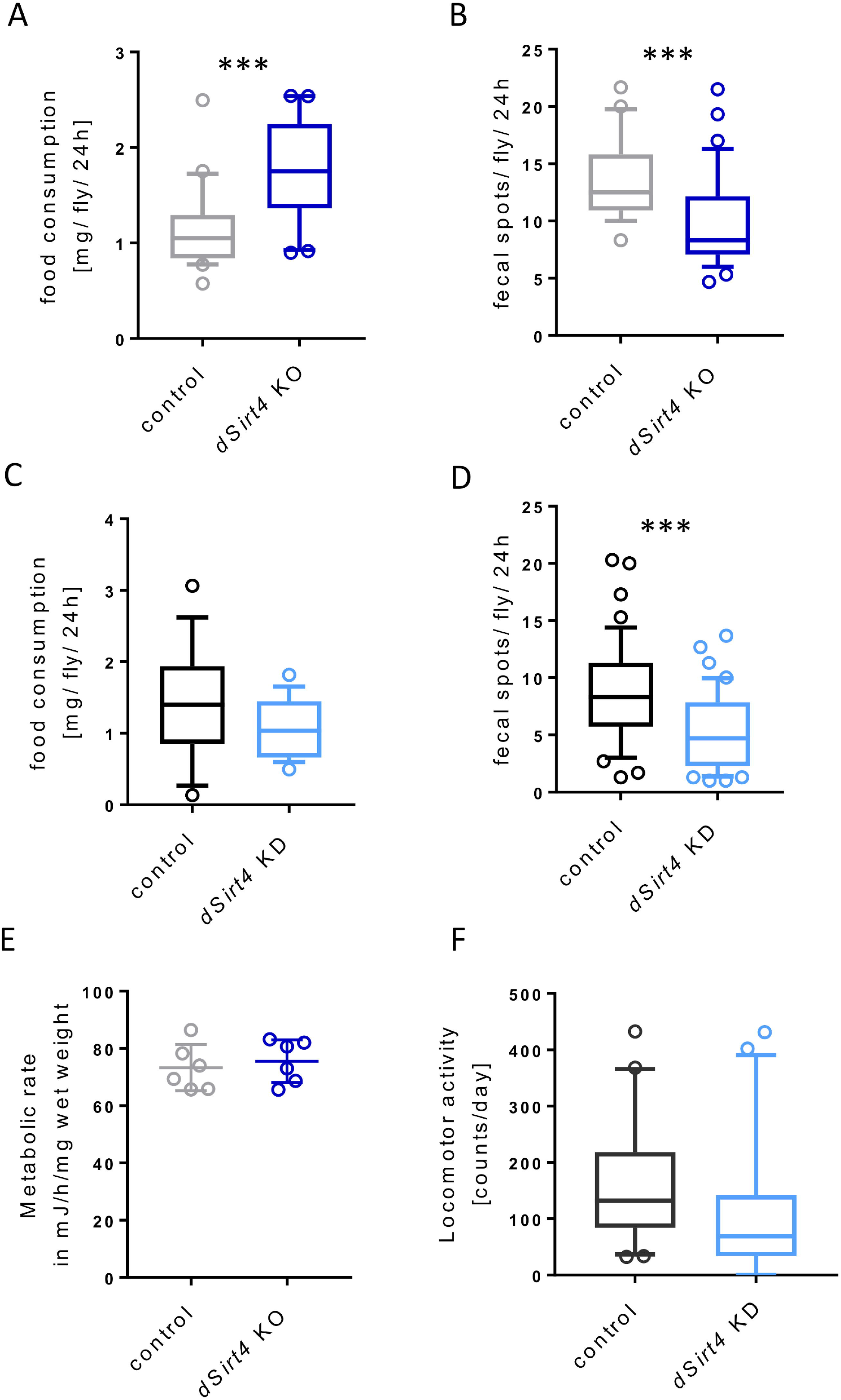
Changes in gut functionality upon reduced *dSirt4* expression. (A, B) The knockout of *dSirt4* (*dSirt4* KO) leads to an increase in food consumption (n = 20), while the number of excreted fecal spots is not affected (n = 26-36). (C, D) In flies deficient of *dSirt4* in enterocytes (*dSirt4* KD), the amount of ingested food is not significantly different compared to the control (n =17-18), but the number of excreted fecal spots is reduced (n = 41). (E) The metabolic rate of *dSirt4* KO flies was determined by measuring the heat dissipation and showed no significant differences (n = 6). (F) Flies with a knockdown of *dSirt4* in enterocytes have a significantly reduced locomotor activity (n = 46-48). (A,B,E) control = *w*^1118^. (C, D, F) control = *w*^1118^ >UAS-sirt4 crispr/Cas9, *dSirt4* KD = NP1-Gal4;tubPGal80ts>UAS-sirt4 crispr/Cas9. *** = p < 0.001.

The effect of a *dSirt4* knockout on the intestinal proteome of *Drosophila* was analyzed by label-free quantitative proteomics using liquid chromatography-mass spectrometry (LC-MS). In 21 analyzed samples, 2881 *Drosophila* protein groups were identified, and 2364 protein groups could be quantified. Between the *dSirt4* KO flies and the control strain, 301 proteins of differential abundance were detected with at least a 2-fold difference and 488 proteins applying a cutoff above 1.5-fold (Fig. 3A, B). The complete dataset is provided in the supplementary Table S1.

**Fig. 3:**
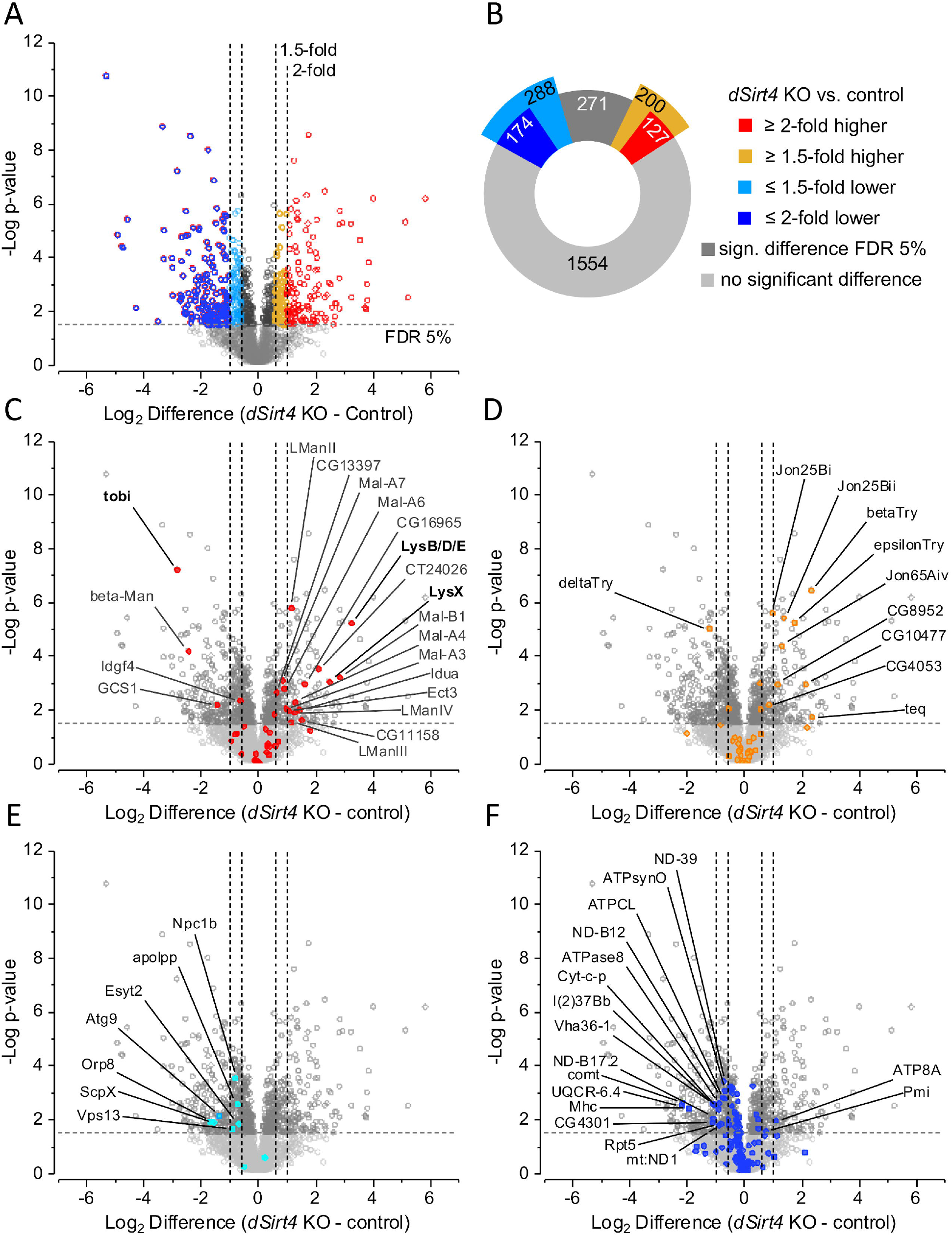
Quantitative proteome analysis. (A, B) Differentially abundant proteins were detected by label-free quantification in *dSirt4* KO intestines (n = 6) compared to the control strain (n = 7) of *Drosophila*. (C) Including the protein group encompassing LysB, LysD and LysE, glycosidases were significantly more abundant, whereas the carbohydrate regulatory protein tobi was less abundant in *dSirt4* KO. (D) Digestive serine proteases were more abundant in *dSirt4* KO intestines. (E) Most proteins involved in lipid transport were of significantly lower abundance in the *dSirt4* KO. (F) Proteins associated with mitochondrial electron transport, the respiratory chain complex, and ATP synthases showed a trend towards lower abundance in *dSirt4* KO intestines. control = *w*^1118^.

Most impressive was the increased abundance of the lysozymes. Due to strong sequence similarity, the lysozymes B, D, and E could only be detected as one protein group, which was of much higher abundance in the intestines of *dSirt4* KO mutants with a 9.5-fold difference compared to the control. In addition, LysX is also found at significantly higher abundances in dSirt4 KO flies (Fig. 3C). Glycosidases were significantly enriched with 16 members among the 200 proteins showing at least a 1.5-fold higher abundance in *dSirt4* KO flies. These included several lysosomal glycosidases for a range of substrates (Fig. 3C). The same trend was observed for proteases. In addition to ten serine-proteases (Fig. 3D), four metalloproteases were more abundant in the *dSirt4* KO flies. In contrast, almost all proteins involved in lipid transport were among the proteins of lower abundance in *dSirt4* KO mutants (Fig. 3E). These data show that even without a change in diet, a *dSirt4* knockout significantly alters the metabolic state of the organism. Interestingly, proteins annotated to the mitochondrial electron transport, the respiratory chain complex, and ATP synthases also showed a remarkable trend towards lower abundance in *dSirt4*-deficient animals compared to the control (Fig. 3F).

To confirm the increased abundance of lysozyme observed in the proteome analysis, we performed a qRT-PCR analysis to measure *lysB* and *lysP* expression. As expected, the expression of *lysB* was significantly up-regulated, however, *lysP* was not (Fig. 4A). Further, we measured the lysozyme activity of dissected intestines by applying the homogenate onto Petri dishes containing agarose mixed with cells walls of *Micrococcus lysideikticus* and quantified the zone of lysis. Intestines dissected from *dSirt4*-deficient flies showed a highly increased lytic activity that was more than 800% higher than that of the control (Fig. 4B), which is in accordance with the difference in protein abundance detected by proteomics. Next, we investigated whether flies with a *dSirt4*-deficiency restricted to enterocytes have a higher lysozyme activity as well. Here, the zone of lysis was also significantly larger, but not to the extent of *dSirt4* KO flies (Fig. 4C). The relative lysozyme activity was increased by approx. 350 %. A qRT-PCR analysis further revealed an up-regulation of *lysB* expression (Fig. 4D). To test if the increased lysozyme activity is part of a broader activation of the immune system, we additionally measured the expression of different antimicrobial peptide genes in the intestine using qRT-PCR analysis. In dSirt4 knockout flies, the expression of *metchnikovin, drosomycin, defensin, diptericin, attacin*, and *cecropin* was significantly down-regulated, only the expression of *drosocin* was not regulated (Fig. 4E). In flies with a dSirt4 deficiency in enterocytes, only the expression of *metchnikovin* was significantly up-regulated, while that of *defensin* was down-regulated (Fig. 4F). All others did not show significant changes in expression.

**Fig. 4:**
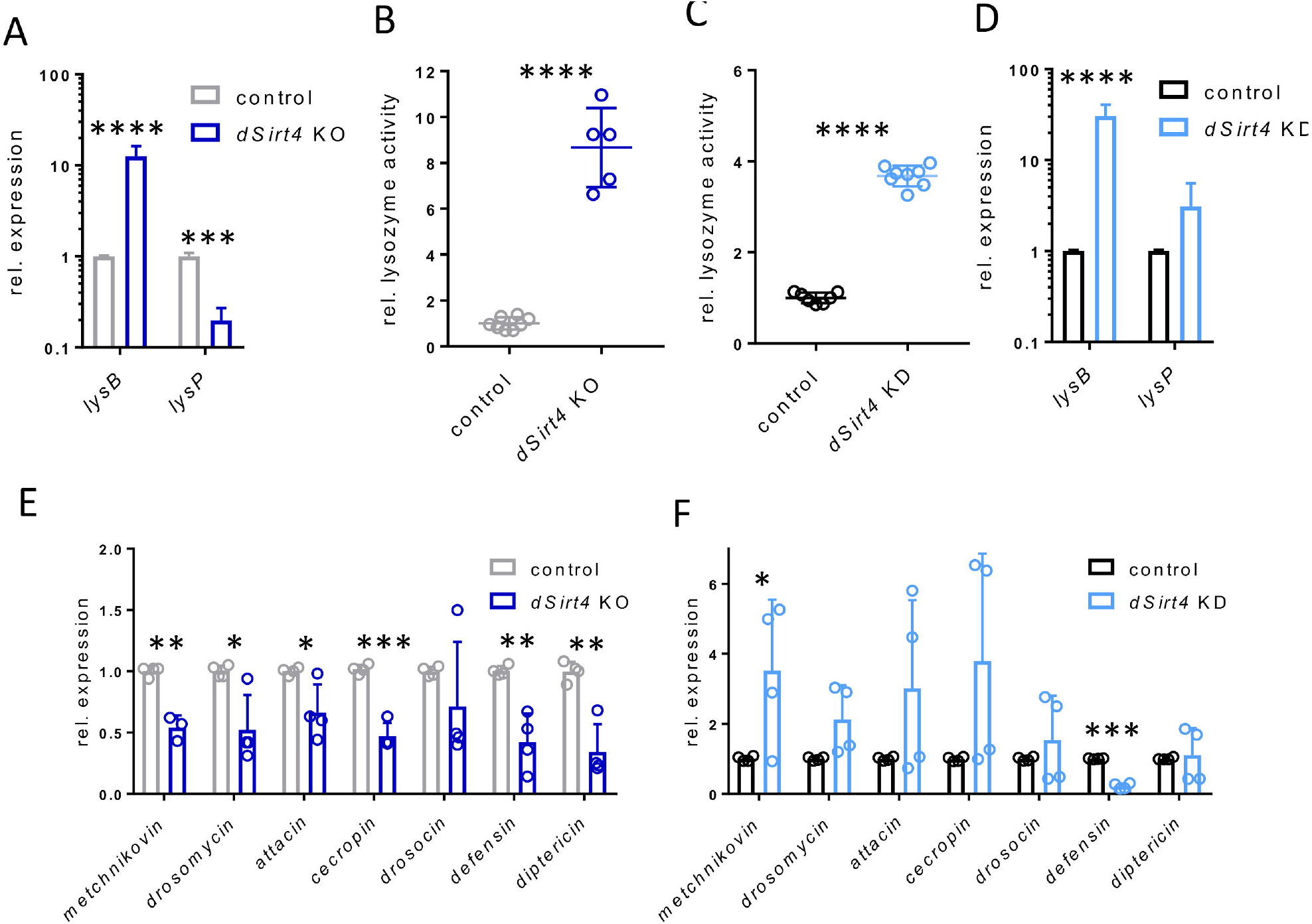
Deficiency of *dSirt4* strongly induces *lys*ozyme expression in the intestine. (A) qRT-PCR analysis showed significantly upregulated *lysB* expression in *dSirt4* KO intestines (n = 4). (B) Strong increase of lysozyme activity in *dSirt4* KO intestines (n = 5-8). (C) Increase of lysozyme activity (n = 8) and (D) upregulation of *lysB* expression in intestines of flies with a *dSirt4* deficiency in enterocytes (n = 4). (E, F) Changes in expression of antimicrobial peptide genes in intestines from *dSirt4* KO flies (E; n =4) and intestines of flies with a *dSirt4* deficiency in enterocytes (F; n = 4). (A,B,E) control = *w*^1118^. (C, D, F) control = *w*^1118^>UAS-sirt4 crispr/Cas9, *dSirt4* KD = NP1-Gal4;tubPGal80ts>UAS-sirt4 crispr/Cas9. * = p < 0.05, ** = p < 0.01, ***= p < 0.001, **** = p < 0.0001.

To test, whether the strong increase in lysozyme activity has an impact on the intestinal microbiota, we performed bacterial load assays with *dSirt4*-deficient, lysozyme-deficient (*lys*^B-PΔ^)and control flies. Axenic embryos were recolonized with a mix of six *Drosophila* gut bacteria (*Lactobacillus brevis, L. plantarum, Acetobacter pomorum, A. thailandicus, Enterococcus faecalis and Commensalibacter intestini*). After hatching, flies were kept together in one container for five days to ensure the same starting condition. Afterwards, flies were sorted and kept for additional ten days. Homogenates of whole flies were plated onto MRS-, LB-, and Mannitol-plates, and the number of colony-forming units (CFU) was determined. The bacterial load of *dSirt4*-deficient flies was significantly increased compared to the control flies in contrast to flies lacking the lysozymes B-P genomic region (Fig. 5A). Nevertheless, due to their ability to cleave β-(1,4)-glycosidic bonds in the peptidoglycan layer and therefore mainly gram-positive bacteria, lysozymes might be involved in altering the microbial composition (33). To test for a shift in the intestinal composition from gram-negative towards gram-positive bacteria caused by the increased activity of lysozymes, flies were treated as mentioned but recolonized with a single gram-positive (*L. plantarum*) and a single gram-negative bacterial species (*A. thailandicus*). Additionally, to the previously used lines, double mutant flies deficient in *dSirt4* and the lysozyme locus (*lys*^B-PΔ^) were used. Compared to the control and to *lys*^B-P^ deficient flies, *dSirt4*-deficient flies had a reduced number of *L. plantarum* (Fig. 5B). The flies lacking *dSirt4* as well as *lys*^B-PΔ^ also showed a higher number of gram-positive bacteria compared to the *dSirt4* deficient flies. The number of gram-negative *A. thailandicus* was not significantly altered in *dSirt4* KO and *lys*^B-PΔ^ flies compared to the control, but slightly reduced in flies double deficient in *dSirt4* and *lys*^B-PΔ^ (Fig. 5B).

**Fig. 5:**
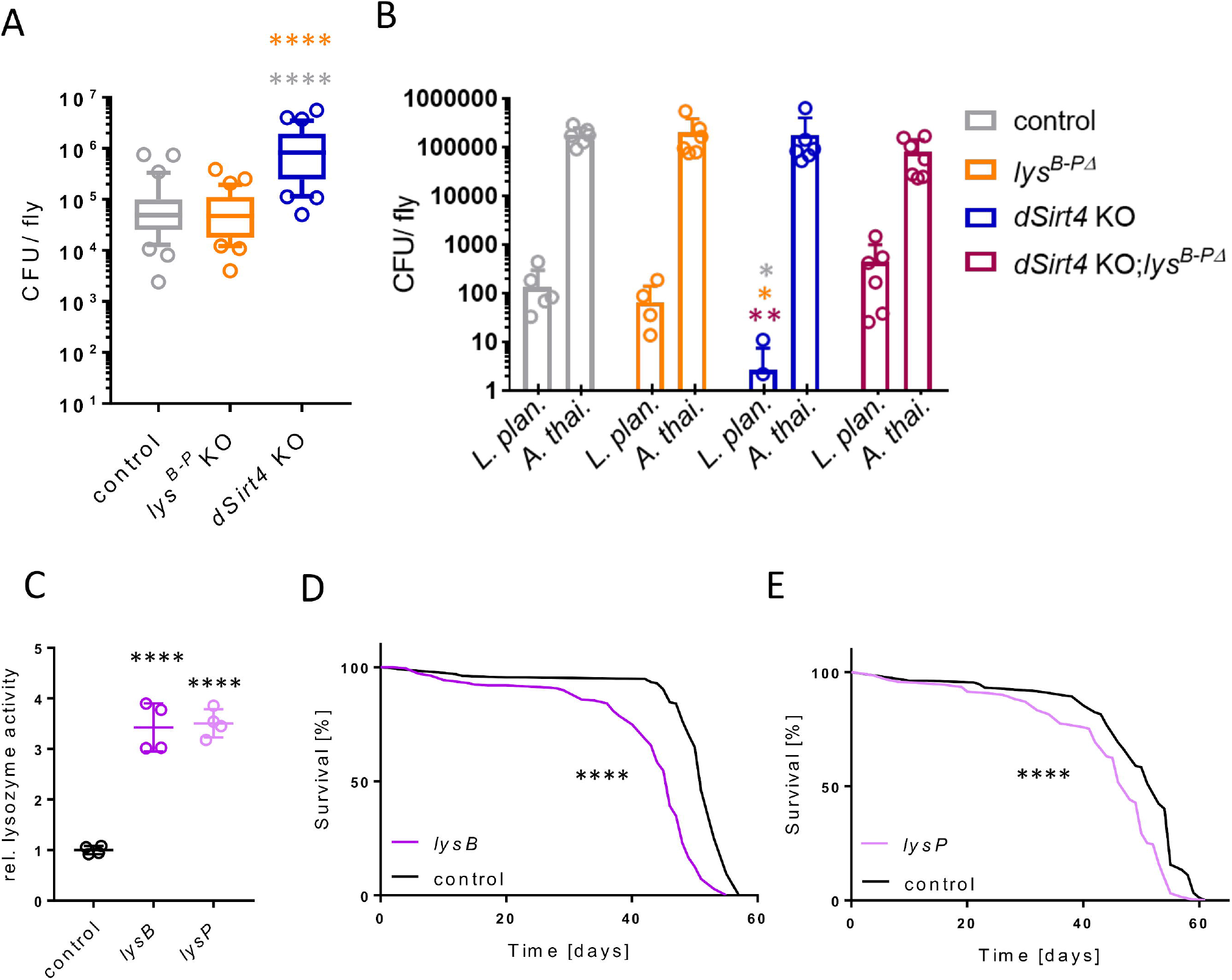
Changes in the bacterial load of intestines upon *dSirt4* KO. (A) Bacterial load of recolonized *dSirt4* K, *lys*^B-PΔ^, and control flies (n = 33). (B) Reduced number of CFU of *L. plantarum* and *A. thailandicus* after recolonization in disassociation *dSirt4* KO flies compared to control, *lys*^B-PΔ^ flies and *dSirt4* KO;*lys*^B-PΔ^ flies (n = 6). (C) Increased lysozyme activity in intestines dissected from flies with overexpressed *lysB* or *lysP* in enterocytes (n = 4). (D, E) Reduced survival in flies with an overexpression of *lysB* in enterocytes (D; n = 150-325) and an overexpression of *lysP* in enterocytes (E; n = 135-314). A, B) control = *w*^1118^. (C) control = NP1-Gal4;tubPGal80ts>*w*^1118^, *lysB* = NP1-Gal4;tubPGal80ts>UAS-*lys*B, *lysP* = NP1-Gal4;tubPGal80ts>UAS-*lys*P. (D) control = *w* ^1118^>UAS-*lys*B, *lysB* = NP1-Gal4;tubPGal80ts>UAS-*lys*. (E) control = *w*^1118^ >UAS-*lys*P, *lysP* = *NP1-Gal4;tubPGal80ts>UAS-lysP*. * = p < 0.05, ** = p < 0.01, **** = p < 0.0001.

Since we found that both the genomic *dSirt4*-deficiency, as well as the knockdown of *dSirt4* in enterocytes, have a highly increased lysozyme activity and both significantly shorten the life span, we addressed the impact of increased lysozyme activity on the survival of *Drosophila*. Therefore, we generated flies carrying either an *UAS-lysB* or an *UAS-lysP* transgenic construct, which were crossed to an enterocyte-specific driver (*NP1-Gal4;tubPGal80ts>UAS-lysB* and *NP1-Gal4;tubPGal80ts>UAS-lysP*). The effectivity of the overexpression was confirmed using a lysozyme activity assay (Fig. 5C). Dissected intestines of flies overexpressing either *lysP* or *lysB* showed a strongly increased activity by approx. 350 % in lysozyme activity. In the survival experiment, flies overexpressing *lysB* in enterocytes lived significantly shorter than the respective control (Fig. 5D). The median life span was reduced by five days (46 days compared to 51 days). The overexpression of *lysP* had the same life-shortening effect (Fig. 5E). Here, the median life span was reduced to 48 days compared to the control with 53 days.

Flies with a genomic *dSirt4*-deficiency are unable to mobilize stored fat in response to starvation (34). Therefore, we were interested in whether the enterocyte-specific deficiency of *dSirt4* shows similar effects. When we measured the body fat content under control? conditions, after 12 hours of starvation, and after 24 hours of starvation, we observed that the triacylglycerol (TAG) content of control flies was significantly reduced in response to starvation (Fig. 6A). *dSirt4*-deficient flies however showed no reduction of their body fat content after a starvation period of 12 hours, indicating flies were not able to mobilize a large part of their fat storage to compensate for the lack of food. In *dSirt*-KD, reduction in body fat was 5 % after 12 h and 28 % after 24 h. The respective reductions for the control were 27 % and 65 %, respectively. We also measured the protein content in response to starvation. Already after 12 hours, the amount of body protein massively decreases (Fig. 6B). After 24 hours, it is even decreased further. The resistance to starvation measured by a life span experiment was also reduced in flies deficient in *dSirt4* in enterocytes (Fig. 6C). The median life span was reduced from 50 to 46 hours compared to the control.

**Fig. 6:**
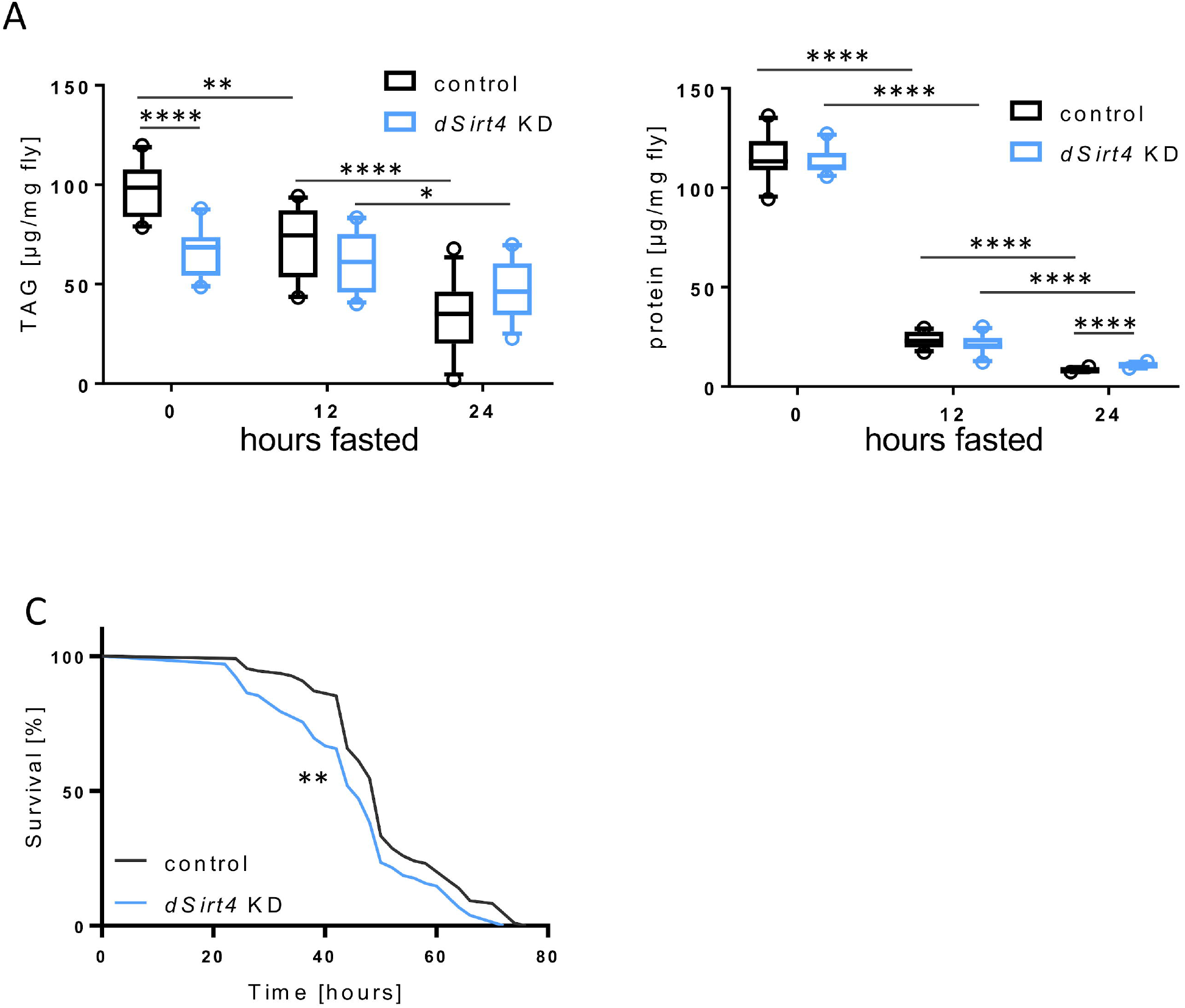
Effects of *dSirt4* on carbohydrate and fat metabolism. (A) Reduced mobilization of fat storages in response to starvation in flies with a knockdown of *dSirt4* in enterocytes (*dSirt4* KD). Bodyfat content was determined by measuring triacylglycerol (TAG) levels (n = 10-12). (B) Massive reduction in protein storage upon starvation in *dSirt4* KD flies over a time period of 24 hours (n = 10-12). (C) Reduced survival of flies with a knockdown of *dSirt4* in enterocytes under starvation condition (n = 102-108). control = *w*^1118^>UAS-sirt4 crispr/Cas9, *dSirt4* KD = NP1-Gal4;tubPGal80ts>UAS-sirt4 crispr/Cas9. * = p < 0.05, ** = p < 0.01, ***= p < 0.001, **** = p < 0.0001.

## Discussion

Building the interface between the environment and the organism, the gut is the organ that is directly confronted with one of the most important environmental factors: nutrition. Information on energy content, but also on the main macronutrients, is of particular interest (35, 36). Adapting to conditions and efficient use of the current diet is therefore one of the main tasks of the gut (37, 38).

To fulfill this task, it needs sensors that enable a profound change in cell metabolism in the short and long term when energy is lacking. Sirtuins are histone deacetylases, which can perform exactly these tasks by acting as indirect energy sensors via NAD+ (9). Following this logic, we looked for those sirtuins that are strongly regulated in the intestine during severe nutritional stress, in our case protein malnutrition. It turned out that only dSirt4, the single mitochondrial sirtuin in *Drosophila*, responds with a substantial up-regulation of its expression. We, therefore, focused on this particular sirtuin and investigated its importance in key aspects of gut biology. It was already known that dSirt4 has a significant influence on life span as overexpression in the fat body has a life-prolonging effect, whereas *dSirt4*-deficient animals live shorter (34).

We showed that *dSirt4*-deficiency, which is restricted to the enterocytes of the intestine, also shortens life span. The comprehensive proteome analysis showed that several proteins showing differential abundances in *dSirt4*-deficient intestines. The massively increased expression of lysozyme was particularly striking. We confirmed this increase at the transcriptome level and the activity level. Again, this effect was observed in the same way when the *dSirt4*-suppression was only suppressed suppressed in enterocytes, showing that the effect is tissue-autonomous and not systemically controlled. Moreover, this dSirt4-dependent increase in lysozyme activity is not part of a complex immune response, as AMP genes were not upregulated under these conditions. However, the mechanism by which decreased dSirt4 activity increases lysozyme activity is still unclear. Lysozyme regulation by mitochondrial sirtuins appears evolutionarily conserved since *lysP* was significantly up-regulated in response to gut-specific knockdown of mitochondrial Sirt3 in mice (15). Lysozymes are known for their antibacterial activity and are therefore generally classified as antibacterial agents. This is due to their ability to lyse gram-positive bacteria by degrading the murein sacculus. Their function as part of the innate immune system against a range of pathogenic bacteria has been demonstrated in various models (39-41). However, recent studies in *Drosophila*, in which the major lysozyme gene loci were deleted, showed that the effect of intestinal lysozymes on the endogenous gut microbiota is not as pronounced as expected (33). We were able to show a similar result, i. e. that increased lysozyme expression did not lead to a massive reduction in bacterial load, but rather to an increase. However, there was a shift in the composition of the microbiota. We chose a very simple model microbiota consisting of two species, the gram-negative *Acetobacter thailandicus* (42) and the gram-positive *Lactobacillus plantarum* (43). The increased lysozyme expression in the *dSirt4*-deficient background had only a minor effect on the microbial load, but significantly changed the composition of this very simple model microbiome. There was a shift towards gram-negative bacteria at the expense of gram-positive bacteria. This shift was due to increased lysozyme expression, as it was not observed in a lysozyme-deficient background. Thus, a direct mechanistic link between the differences in lysozyme expression and the change in the composition of the microbiota can be made.

This complex role of lysozymes in the *Drosophila* intestine can also be observed in the mouse intestine. Here, an imbalance in lysozyme expression in Paneth cells has significant effects on the inflammatory tone of the intestine and thus also on the corresponding chronic IBDs. Lysozyme1-deficiency protects against inflammatory responses, whereas ectopic overexpression promotes inflammatory responses. Thus, excessive lysozyme expression can be correlated with a dysbiotic situation (44). We believe that a similar situation exists in *Drosophila*. This dysbiosis could be the reason for the shortened lifespan of flies with *dSirt4*-deficient enterocytes. This hypothesis is supported by the fact that we were able to show that not only *dSirt4*-deficiency in enterocytes but also the specific overexpression of lysozymes reduces lifespan.

Sirtuin activity is strongly dependent on ingested food and is generally associated with the health-promoting effects of caloric restriction (45, 46). We were able to demonstrate a similar effect in the *Drosophila* gut, but very specifically for dSirt4, while other sirtuin genes tended to be downregulated. Besides nutrition, there are several other factors that may be responsible for the regulation of sirtuin activity. First and foremost is infection with intracellular bacteria of the genus Wolbachia, which is known to lead to a regulation of *dSirt4* expression and impacts the microbiota (26). What effects this has on lysozyme expression and possible dysbiotic composition of the microbiota must be clarified in the future.

In addition to this most interesting result, the increased level of lysozyme and the associated change in the microbiota, *dSirt4*-deficiency leads to other interesting changes in the gut. Of note is the apparent reprogramming of key metabolic properties of the gut, an aspect known to be associated with sirtuins (12, 47). First and foremost are the effects on carbohydrate metabolism and the massive reduction in the expression of tobi, a key regulator of carbohydrate metabolism (48). This is supported by the increased expression of many members of the maltase family, but also of the lysosomal alpha-mannosidases. The increased expression of the latter in the gut is sufficient to induce life span extension (49), a finding that may be very interesting for the interpretation of our results. Furthermore, we found a reduced abundance of proteins that are operative in lipid transport processes that might be a reason for the reduced ability to utilize fat reserves in response to starvation. Here, the reduced abundance of aplopp and Npc1b might be of greatest importance (50, 51). In summary, we have identified dSirt4 as a highly sensitive cellular sensor in the adult *Drosophila* gut that not only responds to changes in diet, but also broadly reprograms gut metabolism. Of particular note is the expression of lysozyme, which provides a direct mechanistic link to the composition of the microbiota and thus significantly advances our understanding of the development of chronic inflammatory diseases of the gut, where sirtuins might play an important role (17).

## Material and Methods

### Drosophila Stocks and Culture

The following strains were obtained from the Bloomington Stock Center: *w*^1118^ (#5905), *dSirt4* KO (#8840), UAS-Cas9 (#54592) and *Sirt4* sgRNA (#78741). Sirt4 KO flies were backcrossed to w^1118^ flies for several generations prior experiments. NP1-Gal4; tubPGal80ts was kindly provided by D. Ferrandon and Lys^B-PΔ^ by Bruno Lemaitre. UAS-sirt4 crispr/Cas9 was generated by combination of UAS-Cas9 (#54592) and *Sirt4* sgRNA (#78741). *UAS-lysB* and *UAS-lysP* were generated using the pBID-UASC vector (REF?) and EcoRI, BglII and BamHI restriction enzymes. cDNA of *lys*B-PA was amplified with GAGAATTCCAAAATGAAGGCTTTCATCGTTCTG and GAGGATCCGAAGCAGTCATCGATGGAC primers, *lysP* was amplified with GAGAATTCCAAAATGAAAGCTTTTCTTGTGA and GAAGATCTGCAACTGTTGATCGAGGGCA primers. Flies were cultivated on a standard diet (NM, per 500 ml: 31.25 g brewer’s yeast, 31.25 g cornmeal, 10 g D-glucose, monohydrate, 5 g agar agar, 15 g sugar beet syrup, 15 g molasses, 5 ml of propionic acid (10 % in ddH_2_O) and 15 ml of nipagin (10 % in 70 % EtOH) for preservation). Temperature sensitive crossings were raised at 18°C, all others at 25°C. If not stated otherwise, five to seven day old female flies were used for experiments. The F1 progeny containing temperature-inducible genetic modules and their corresponding controls were kept at 18°C for five days before the induction of the expression of the gene of interest at 29°C for five days. For applying protein malnutrition, a holidic diet was used as described earlier (28).

### Life span and infection survival time

The life span experiment was performed in standard *Drosophila* vials. Flies (females) were monitored daily and transferred onto fresh NM every two to three days until all flies had died.

The life span of flies under starvation condition was performed in standard *Drosophila* vials filled with approx. 10 ml of 1.5 % agar-agar to prevent a dying of thirst. Dead flies were counted every two hours until all flies had deceased. The influence of DSS (MP Biochemicals, Canada) on the life span of *Drosophila* was tested in standard *Drosophila* vials filled with approx. 10 ml of 1.5 % agar-agar. A solution of 5 % DSS (w/v) in 5 % sucrose (w/v) was applied on stripes of filter paper. The filter paper was exchanged every two days, the vials once a week. Dead and escaped flies were counted every day (sucrose controls every second day).

For the survival upon infection, overnight cultures of Serratia marcescens (Db11) were grown at 30°C in lysogenic broth (LB) medium supplemented with Streptomycin (10 μg/ml). Cultures were concentrated by centrifugation and resuspended in 5 % sucrose to OD_600_ = 50. The bacterial solution was applied on stripes of filter paper. The filter paper was exchanged every two days. Controls were fed with 5% sucrose only.

### Food Consumption Assay

The consumption of food was assessed with the previously described consumption-excretion method (Shell et al. 2018) with minor adjustments. NM or blue dyed NM (0.5 % (w/v) Brilliant Blue FCF food dye; E133) was pipetted into caps of 2 ml screw cap vials. Individual flies were transferred into 2 ml screw cap vials with regular NM. Vials were loosely closed to ensure air supply. After several hours of adaptation, caps were replaced with blue NM. After 24 hours caps containing blue NM were replaced with clean empty ones. Flies were homogenized with 500 μl H_2_O using a bead ruptor (OMNI International, Kennesaw, USA). The homogenate was centrifuged at 3000 rpm for 3 min to pellet the tissue debris and transferred into 96-well plates. The absorbance at 630 nm was quantified using a SYNERGY H1 microplate reader (BioTek Instruments, Vermont, USA). A standard curve of a dilution series of the blue food was used to calculate the amount of ingested food.

### Measurement of fecal output

Standard *Drosophila* vials were placed tilted and filled with approx. 3 ml of blue dyed NM (0.5 % (w/v) Brilliant Blue FCF food dye and a cover slip (24 × 50 mm). Three female flies were placed in each experimental vial. The cover slip was fixed with the plug and served as the bottom. After 24 hours, all fecal spots were counted and calculated per fly.

### Body composition

The body fat of *Drosophila* was measured using the coupled colorimetric assay (CCA) from (52) as described by (53). The protein content was determined using the Pierce™ BCA Protein Assay Kit (ThermoFisher, Karlsruhe, Germany) according to the manufacturer’s protocol.

### Activity monitoring

The activity of flies was measured using the *Drosophila* Activity Monitor System (DAM, TriKinetics. Waltham, USA) as previously described with minor modifications (54). Individual flies were transferred into glass tubes filled with NM. Tubes were placed horizontally into the DAM device. After adaptation to the conditions for one day, the activity was monitored for three additionally days. The data was analyzed using the web application ShinyR-DAM (55).

### Metabolic rate

The metabolic rate of individual adult female flies was determined by direct microcalorimetry in a TAM IV instrument equipped with six 4ml microcalorimeters (TA Instruments) as described (56). In detail, *dSirt4* heterozygous mutant flies were backcrossed to *w*^1118^ (BDSC #5905) flies for three generations before *w*^1118^ TI{*w*^+mW.hs^=TI}Sirt4^white+1^ / *w*^1118^ and *w*^1118^ virgin females were crossed to *w*^1118^ TI{*w*^+mW.hs^=TI}Sirt4^white+1^ / Y and *w*^1118^ males, respectively. The fertilized females were pooled in the same NM vial and homozygous *Sirt4* mutant and *w*^1118^ control F1 females were mated and afterwards kept separately from the males at 25°C in constant darkness and at 75% relative humidity for six days on NM. Individual flies were transferred to disposable 4ml crimp seal glass ampoules preloaded with 200μl NM. Heat dissipation of a total of six flies per genotype (two runs of three mutants and controls each at two consecutive days) was measured over four hours. After the microcalorimeter run, the wet weight of the flies was determined using a Sartorius MC5 balance. Heat dissipation of individual flies was averaged per hour and the metabolic rate was calculated in mJ/h/mg wet weight.

### Sample preparation for proteome analysis

*Drosophila* intestines of *dSirt4*-KO flies and the control strain (*w*^1118^) were analyzed by bottom-up proteomics and label-free quantification. *Drosophila* intestines were prepared as described previously (29).

Cell disruption was performed in 5 μL of lysis buffer (6M Urea, 100 mM TEAB, 1x complete protease inhibitor) per *Drosophila* intestine together with glass beads. First samples were processed using a Bioruptor pico with ten cycles of 30s of sonication and 30s of cooling followed by vortexing of 10 s and freezing at -80°C. These disruption steps were repeated five times and the protein concentration was determined by BCA assay. Aliquots of 20 μg protein were reduced for 1h at 60°C with 20 mM Tris (2-carboxyethyl) phosphine (TCEP) and then alkylated for 30 min with 40 mM Chloroacetamide (CAA) and digested overnight with 0.5 μg of trypsin. After digestion, the samples were acidified to 0,1% TFA, lyophilized to dryness, resolved in 100 μL 0.1% TFA, and clean-up using 100 μL C18-tips according to the protocol of the manufacturer (Pierce). The samples were again lyophilized to dryness, dissolved in 20 μL 0.1% TFA, and transferred to vials for HPLC separation.

### LC-MS analysis

For LC-MS analysis, approximately 1 μg of peptides were loaded on a C18 precolumn (PepMap100, 5 μm, 300 Å, Thermo Fisher Scientific) and separated over a C18 column (50 cm × 75 μm, 2.6 μm, 100 Å, Thermo Fisher Scientific) using a Dionex U3000 UHPLC system (Thermo Fisher Scientific) coupled to a Q Exactive HF mass spectrometer (Thermo Fisher Scientific). The separation was performed across a 2.5 h gradient with eluent A (Water, 0.05 % FA) and eluent B (80% ACN, 0.04 FA) with a flow rate of 0.3 μL/min. First for 5 min at 5% B; in 80 min to 20% B; in 65 min to 50% B; in 5 min to 90% B; isocratic at 90% B for 10 min and equilibration for 10 min at 5% B.

Full MS spectra were acquired with the following settings: resolution of 601000, mass range of 300–1600 m/z, 30% RF lens, AGC target of 3E6, and 100 ms maximum injection time. MS2 spectra of the top 10 precursors with a charge state above 2 and below 8 were acquired with an isolation window of 2 m/z, 151000 resolution, AGC target of 2E5, 100 ms injection time, and an NCE of 28. Dynamic exclusion was enabled with an exclusion duration of 10 s.

In total, 21 samples were analyzed and a database search and label-free quantification were executed with Proteome Discoverer software (version 3.0.1.27; Thermo Fisher Scientific). The raw data were searched against the combined UniProt protein databases of *Drosophila melanogaster* (UniProt 05.2023; 22,066 entries), the defined microbiota bacteria: *Lactobacillus plantarum* (UniProt 07.2021; 3,179 entries); *Lactobacillus brevis* (UniProt 07.2021; 2,201 entries); *Acetobacter pomorum* (UniProt 07.2021; 2,815 entries); *Commensalibacter intestine* (UniProt 07.2021; 2,209 entries); *Enterococcus faecalis* (UniProt 07.2021; 3,240 entries), and common contaminants. Sequest HT and Chimerys search algorithms were used. The Sequest search parameters were semi-tryptic protease specificity with a maximum of 4 missed cleavage sites. The precursor mass tolerance was 10 ppm, fragment mass tolerance was 0.04 Da. Oxidation of methionine and acetylation of lysine residues and protein N-termini were allowed as dynamic modifications. Carbamidomethylation of cysteine was set as static modification. The default settings were used for Chimerys database. Percolator q-values were used to restrict the false discovery rate (FDR) of peptide spectrum matches to 0.01. The FDR of peptide and protein identifications was restricted to 1% and strict parsimony principles were applied to protein grouping. Label-free quantification was performed with Minora feature detector. Label-free intensities were based on the precursor intensities of MS spectra and unique and razor peptides were used to calculate label-free intensity values (LFI) of protein groups.

Statistical analysis was performed with the Perseus software. The dataset was filtered for 2313 protein groups with at least six valid label-free intensity values (LFI). Label-free intensities were log_2_ transformed and missing values were replaced from a normal distribution (width: 0.3, downshift: 2). The datasets of *dSirt4* KO and control samples grown on defined microbiota with and without imputation were tested for differentially abundant proteins using Welch’s t-test and corrected for multiple testing by permutation-based FDR analysis with 250 randomizations. Proteins with a q-value below 0.05 and a 2-fold or 1.5-fold difference were set as two classifications of differential abundance. Fisher’s exact test was used to analyze the annotation enrichment of GO terms and UniProt Keywords in Proteins categorized as higher and lower abundant in *Sirt4* KO mutants. The annotation enrichment analysis was corrected for multiple testing using Benjamini Hochberg FDR calculation.

The results of the proteome analysis are provided in the supplementary table S1. MS data were deposited to the ProteomeXchange Consortium (57) by the PRIDE partner repository with the dataset identifier PXDXXXXX.

### Lysozyme assay

To measure the lysozyme activity, homogenates of *Drosophila* intestines were pipetted onto agarose plates containing cell walls of *Micrococcus lysodeikticus*. For plates, 0.05 M NaAC was mixed with 0.9 % agarose and boiled. 0.6 mg/ml *Micrococcus lysodeikticus* ATCC No. 4698 (Sigma Aldrich, M3770-5g) was solved in 1 ml NaAc at 37°C and shaking. After the agarose cooled down to under 50°C, the *Micrococcus* solution was added and poured into petri dishes. 5 intestines were dissected and homogenized in 50 μl PBS using a bead ruptor (OMNI International, OMNI Bead Ruptor 24). The homogenate was added into the holes punched into plates. After incubation at 37°C for 24h, the diameter of the lysis zone was measured.

### Bacterial load assay

For egg deposition, flies were placed on apple juice agar and several chunks of fresh yeast mixed with few drops of apple vinegar. After 18 h at 20 °C, eggs were dechorionated with 6 % NAClO for five minutes, sprayed with 70 % EtOH, rinsed with sterile water and placed onto sterile NM without propionic acid. The germfree embryos were recolonized with a mix of six bacteria species: *Lactobacillus plantarum*^WJL^, *Lactobacillus brevis*^EW^, *Acetobacter pomorum, Commensalibacter intestini*^A9111T^ *and Enterococcus faecalis* which were all received as a kind gift from Carlos Ribeiro and *Acetobacter thailandicus*, a gift from Luis Teixeira. The culturing and the adjustment of specific optic densities were performed as described by Santos et al. (2017). Each *Drosophila* vial was inoculated with 50 μl of the bacterial suspension. For the association with only two bacterial strains, embryos were recolonized with 50 μl of OD_600_ = 2 of *A. thailandicus* and *L. plantarum* each. After hatching, flies were inoculated with fresh bacteria solution and transferred to sterile NM every three to four days. On day 10, groups of three flies were homogenized in 100 μl sterile PBS. Five dilutions (up to 1:10000) were plated on MRS, LB and Mannitol-Agar plates (58).

### Total RNA Extraction

Total RNA was extracted using RNA Magic and the Ambion PureLink™ RNA Mini Kit (Thermo Fisher, Karlsruhe, Germany). Eight to ten dissected *Drosophila* intestines homogenized in 1 ml RNAmagic (Bio-Budget Technologies GmbH, Krefeld, Germany) using the bead ruptor (OMNI International, Kennesaw, USA). After incubation at RT for 5 min, 200 μl of chloroform was added. Samples were shaken for 10 s, incubated on ice for 5 min and centrifuged at 4°C and 12,000 x g for 15 min for phase separation. 400 μl of the upper phase containing the RNA was transferred into a 1.5 ml reaction tube. The RNA was purified according to the manufacturer’s protocol (Purifying RNA from Animal Tissue; steps Binding, Washing; and Elution). Samples were eluted in 30 μl RNase-free water and stored at -80°C. Reverse transcription of mRNA was performed to generate cDNA using SuperScriptIV reverse transcriptase (Thermo Fisher, Karlsruhe, Germany).

### Quantitative real time PCR

Quantitative real time PCR was performed using the qPCRBIO SyGreen Mix Hi-Rox (PCR Biosystems, London, UK), MicroAmp Optical 96-well Reaction plates (0.1 ml, Thermo Fisher, Karlsruhe, Germany) and the QuantStudio 1 Real-Time PCR System (Thermo Fisher, Karlsruhe, Germany). Following primer were used: *drosomycin* (ACCAAGCTCCGTGAGAACCTT and TTGTATCTTCCGGACAGGCAG), *metchnikowin* (CCACCGAGCTAAGATGCAA and AATAAATTGGACCCGGTCTTG), *diptericin* (GCAATCGCTTCTACTTTGGC and TAGGTGCTTCCCACTTTCCA), *attacinA* (TTCCGTGAGATCCAAAG and CAATCTGGATGCCAAGGTCT) *cecropin* (AAGATCTTCGTTTTCGTCGC and GTTGCGCAATTCCCAGTC), *drosocin* (GTTCACCATCGTTTTCCTGC and GGCAGCTTGAGTCAGGTGAT), *defensin* (GCTATCGCTTTTGCTCTGCT and GCCGCCTTTGAACCCCTTGG), *lysP* (CCAGGCCCGAACGATGGATAGGT and CGGGGAACGCCCAGTTTGGA), *rpl32* (CCGCTTCAAGGGACAGTATC and GACAATCTCCTTGCGCTTCT).

## Statistical analysis

Statistical analysis for life span assay was done using Log-rank (Mantel-Cox) test. All other data was tested for normal Gaussian distribution by Shapiro-Wilk normality test and then accordingly further analyzed using Unpaired t-test or Unpaired t-test with Welch’s correction.

## Acknowledgments

This work was supported by CRCs 1182 (TPC2, A2, Z3), funded by the Deutsche Forschungsgemeinschaft (DFG), Germany; the Clusters of Excellence ‘‘Inflammation@interfaces’’ and ‘‘PMI’’; and the Leibniz Science Campus Evolung; the University of Graz (to R.K.).We would like to thank Dominique Ferrandon, Bruno Lemaitre, and the Bloomington Stock Center for flies. Moreover, we would like to thank Britta Laubenstein, Christiane Sandberg, Lisa Pichler and Lydia Misslinger for excellent technical assistance.

## Author Contributions

Experimental design/discussion, M.K., J.B., R.K., J.v.F., I.B., J.F.B. A.T. and T.R.; preparation and performance of experiments and data analysis, M.K., C.T., S.B., J.B., J.v.F. R.K. and S.D.; manuscript preparation, M.K., R.K., A.T. and T.R. All authors agreed on the contents, including the author list and author contribution statements.

